# A new symbiotic lineage related to *Neisseria* and *Snodgrassella* arises from the dynamic and diverse microbiomes in sucking lice

**DOI:** 10.1101/867275

**Authors:** Jana Říhová, Giampiero Batani, Sonia M. Rodríguez-Ruano, Jana Martinů, Eva Nováková, Václav Hypša

## Abstract

Phylogenetic diversity of symbiotic bacteria in sucking lice suggests that lice have experienced a complex history of symbiont acquisition, loss, and replacement during their evolution. By combining metagenomics and amplicon screening across several populations of two louse genera (*Polyplax* and *Hoplopleura*) we describe a novel louse symbiont lineage related to *Neisseria* and *Snodgrassella*, and show its’ independent origin within dynamic lice microbiomes. While the genomes of these symbionts are highly similar in both lice genera, their respective distributions and status within lice microbiomes indicate that they have different functions and history. In *Hoplopleura acanthopus*, the *Neisseria*-related bacterium is a dominant obligate symbiont universally present across several host’s populations, and seems to be replacing a presumably older and more degenerated obligate symbiont. In contrast, the *Polyplax* microbiomes are dominated by the obligate symbiont *Legionella polyplacis*, with the *Neisseria*-related bacterium co-occurring only in some samples and with much lower abundance.

## Introduction

An increasing number of studies demonstrate ubiquity and high diversity of insect-associated microbiomes (Douglas 2015, Engel and Moran 2013). These microbial communities, composed of various pathogens, commensals and random contaminants, can serve as natural sources of beneficial symbiotic bacteria. In some insects they give rise to highly specialized, maternally-transmitted mutualists called primary symbionts (P-symbionts), which contribute to the host’s metabolism (Douglas 1989). However, depending on richness and dynamics, the microbiomes usually contain several symbiotic bacteria in various evolutionary stages. In their typical form, P-symbionts are readily recognized by several features (since they are indispensable mutualists): they are universally present in all individuals, as a rule inhabiting specialized host’s organs called bacteriomes (Baumann 2005), and their genomes are significantly reduced with a strong AT bias (Moran 1996). One specific feature of P-symbionts is their co-phylogeny with the host (Chen et al 1999, Clark et al 2000, Sauer et al 2000). For example, two of the most studied P-symbionts, *Buchnera* in aphids and *Wigglesworthia* in tsetse flies, were acquired at the beginning of their hosts’ diversification and strictly mirror their entire phylogeny (Chen et al 1999, Clark et al 2000). Other P-symbionts are restricted to some of the host’s lineages, indicating that they are either recently acquired symbionts or remnants of an ancient symbiont lost in some of the host lineages (Bennett and Moran 2013). In some insects, several different P-symbionts may coexist and/or can be accompanied by various secondary symbionts (S-symbionts). The latter are less modified, retain more free-living-like characteristics, and some are supposed to be the intermediate stages of evolution towards obligate symbionts. *Wigglesworthia* represents a typical example of this as it is often accompanied by the S-symbionts *Sodalis glossinidius* and *Wolbachia* (Aksoy 2000). The complexity of symbiotic associations are obviously due to an ongoing process of symbiont acquisition/loss/replacement, which is well known from several bacteria-insect models and has a well-developed theoretical background (Bennett and Moran 2015). The theoretical work postulates that after a certain amount of coevolutionary time, the symbiotic bacterium becomes too degenerated and functionally inadequate, and it has to be replaced (or accompanied) by another symbiont. While it would be interesting to see how the microbiome diversity and dynamics relate to the complexity of symbiosis in different insect groups, there is very little information available today. The majority of studies on insects and their P- and S-symbionts relies on metagenomic information and phylogenetic reconstructions, likely missing a substantial part of microbiome diversity. The introduction of amplicon approaches recently demonstrated that this method can significantly improve our insight into microbiome composition, even in extensively studied model systems (Doudoumis et al 2017, Gauthier et al 2015, Manzano-Marin et al 2017, Meseguer et al 2017).

Amongst hematophagous (blood-feeding) insects which live exclusively on vertebrate blood, sucking lice of the order Anoplura, with more than 500 spp. (Light et al 2010), are the most ancient and diversified group. Accordingly, they possess a high diversity of symbiotic bacteria (Allen et al 2016, Boyd et al 2014, Boyd et al 2016, Fukatsu et al 2009, Hypsa and Krizek 2007). Depending on interpretation, the 16S rRNA gene-based phylogenies for the available taxa suggest 5-6 independent symbiotic lineages. However, none of them is a universal louse symbiont distributed across the whole order (e.g. like *Buchnera* in aphids). The distribution of louse symbionts suggests a relatively recent origin of each lineage and hence a high rate of acquisition/loss/replacement processes. Moreover, compared to the extensively screened phytophagous groups, only a small fraction of sucking lice diversity has been investigated. The actual number of symbiotic lineages is therefore likely to be much higher. Of the currently known lineages, genomic data are only available for four; three of them showing clear signatures of P-symbionts: *Riesia* spp., *Puchtella pedicinophila*, and *Legionella polyplacis* (Table 1). Correspondingly, each of these lineages has been found in two to four related host species as a result of co-phylogenetic processes. The fourth lineage, the *Sodalis*-like symbiont from *Proechinophthirus fluctus*, possesses a significantly larger genome exceeding 2 Mbp, and GC content 50%, which the authors interpret as possible evidence of recent replacement of a more ancient and now extinct endosymbiont (Boyd et al 2016). The diversity and distribution of the known symbionts in sucking lice thus indicate that this insect group has been undergoing particularly dynamic acquisition, loss, and replacement of symbionts. In this study, we analyze the background of these processes by combining genomic and amplicon approaches across several populations of the louse genera *Polyplax* and *Hoplopleura*. We reveal a new symbiotic lineage related to the genera *Neisseria* and *Snodgrassella* (the latter being a symbiont of bees). We show that these bacteria established their symbiotic relationships independently with the two louse genera, and we prove their intracellular localization in host’s bacteriocytes. Based on the phylogeny-dependent diversity of the microbiome profiles, we suggest rapid microbiome changes at the host population level, possibly underlying the dynamic processes of symbiont acquisition, loss, and replacement in these blood sucking insects.

**Table 1.**
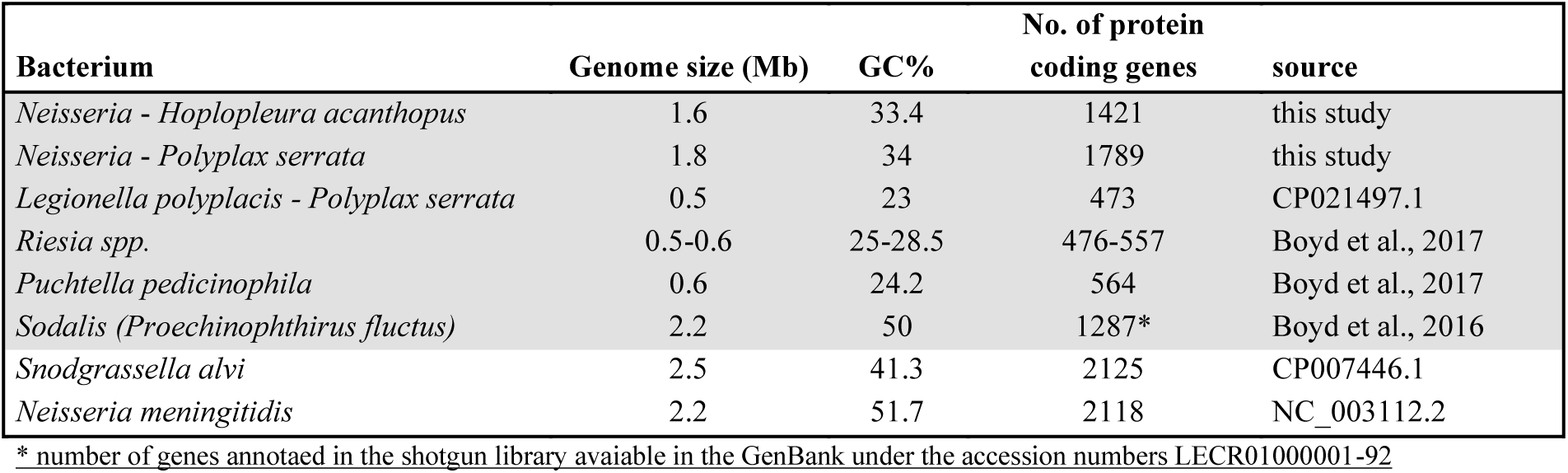
Comparison of main genomic characteristics. Louse symbionts highlighted by grey background.

| Bacterium | Genome size (Mb) | GC% | No. of protein coding genes | source |
| --- | --- | --- | --- | --- |
| *Neisseria - Hoplopleura acanthopus* | 1.6 | 33.4 | 1421 | this study |
| *Neisseria - Polyplax serrata* | 1.8 | 34 | 1789 | this study |
| *Legionella polyplacis - Polyplax serrata* | 0.5 | 23 | 473 | CP021497.1 |
| *Riesia spp.* | 0.5-0.6 | 25-28.5 | 476-557 | Boyd et al., 2017 |
| *Puchtella pedicynophila* | 0.6 | 24.2 | 564 | Boyd et al., 2017 |
| *Sodalis (Proechinophthirus fluctus)* | 2.2 | 50 | 1287* | Boyd et al., 2016 |
| *Snodgrassella alvi* | 2.5 | 41.3 | 2125 | CP007446.1 |
| *Neisseria meningitidis* | 2.2 | 51.7 | 2118 | NC_003112.2 |

\* number of genes annotaed in the shotgun library avaiable in the GenBank under the accession numbers LECR01000001-92

## Methods

### DNA template preparation

*Polyplax serrata* lice (n=25) were collected from yellow-necked wood mice (*Apodemus flavicollis)* captured in Czech Republic (Struzna) and Germany (Baiersbronn) during 2011. *Hoplopleura acanthopus* lice (n=40) were obtained from common voles (*Microtus arvalis)* trapped in Czech Republic (Hlinsko) during 2014. All samples were stored in 96% ethanol at −20°C. Total DNA was extracted from whole louse abdomens using QIAamp DNA Micro Kit (Qiagen) according to manufacturer protocol. DNA quality was assessed by gel electrophoresis and its’ concentration measured with Qubit High sensitivity kit.

### Genome sequencing and assembly

We sequenced the *Polyplax* lice pooled sample on one lane of Illumina HiSeq2000 (GeneCore, Heidelberg) using 2×100 paired-end reads. Read quality was checked using FastQC (Andrews 2010) and quality trimming was performed using the BBtools package (https://jgi.doe.gov/data-and-tools/bbtools). The resulting dataset contained 309,892,186 reads. We used SPAdes assembler 3.10 (Bankevich et al 2012) to build the assembly, implementing careful options and enabling mismatch corrections. To check for bacterial plasmid(s) we submitted the complete assembly (124,985 contigs) to the PlasmidFinder (Carattoli et al 2014) with sensitivity set to three different thresholds (95%, 85%, and 60%). We identified bacterial contigs by blasting *Snodgrassella alvi* wkB2 genome against the assembly using custom blast in the program Geneious (Kearse et al 2012). This procedure retrieved 39 contigs which were putatively assigned to Neisseriales and their origin was further confirmed by blast analyses of individual genes as specified below.

To sequence a complete *Hoplopleura acanthopus* lice metagenome, we employed Illumina MiSeq (GeneCore, Heidelberg) and Oxford-Nanopore (University of Urbana, Illinois) technology. We constructed the Illumina library from the total DNA of 35 individuals and sequenced it in four runs of Illumina MiSeq using V2 500 cycle chemistry. We used the same procedure for quality checking and filtering as described for the *Polyplax* data set. The resulting number of reads was 34,406,078. We used high molecular weight DNA from 5 *H. acanthopus* as a template for Oxford-Nanopore sequencing on GridIONx5. The total number of reads was 1,653,194. The quality of Nanopore reads was checked using NanoPack tools (De Coster et al 2018) and quality filtering was performed using Filtlong (https://github.com/rrwick/Filtlong).

To assemble the *H. acanthopus* metagenome a hybrid approach combining the Illumina and Nanopore data was employed. We used two assemblers, Flye (Kolmogorov et al 2019) and Canu (Koren et al 2017), to generate contigs from Nanopore reads. While Flye assembly resulted in 724 contigs, Canu assembler generated 2,762 contigs. The Nanopore filtered reads were mapped back on both assemblies using Minimap2 (Li 2018). To polish the contigs subsets we used consensus calling in Racon (Vaser et al 2017) followed by two iterations of Medaka polish (https://github.com/nanoporetech/medaka). To obtain optimum sequence correctness, the resulting contigs of the two assemblers were polished with Illumina trimmed reads using Minimap2 alignment and Racon contigs consensus polish. Corrected Flye and Canu assemblies consisted of 702 and 2,721 contigs, respectively. We identified three bacterial contigs using the same blast procedure as described above for *P. serrata*, and assembled them into a single linear sequence (with more than 500 bp congruent overlaps) using the De Novo Assembly tool in Geneious. The genome was completed and closed with an additional 1,643 bp contig retrieved from the corrected Canu assembly into the 1,607,498 bp long circular genome.

We annotated the genomes of both *Neisseria*-related symbionts using RAST (Aziz et al 2008) and deposited them in GenBank under the accession numbers CP046107 (closed genome of the symbiont from *H. acanthopus*) and WNLJ00000000 (draft genome of *Polyplax serrata* symbiont in 39 contigs). To assess the phylogenetic origin of individual genes, we first blasted the complete set of protein-coding genes against the non-redundant (nr) protein database using the blastp algorithm (Altschul et al 1990) set to retrieve one hit. The genes which returned members of Neisseriales were assigned to a category “Neisseriales” (Supplementary Data 1) and considered to be inherited from the Neisseriales ancestor. The rest of the genes were blasted again with blast parameters set to ten hits. Based on the results, the genes were assigned to the following categories: “Mixed” if the hits contained any Neisseriales together with other bacterial groups, “Putative HGT” if the hits did not contain any of the Neisseriales (or even any of betaproteobacteria; designated by bold **Putative HGT** in the Supplementary Data 1), “E” if the hits were eukaryotic, and “No hit” if not hit was obtained. Two bacteria were removed from the nr database prior to this analysis, *Francisella* sp. (accession number GCA_003248485.1) and *Haemophilus parainfluenzae* (GCA_003240835.1). During our preliminary analysis, genes of these two bacteria formed a substantial part of the best hits. Upon closer inspection, we found these two bacteria misclassified (most likely being Neisseriaceae or perhaps chimeras obtained from environmental metagenomic samples).

### Genome comparison

Genome synteny was analyzed using ProgressiveMauve (Darling et al 2010). The complete closed genome of the *Neisseria*-related symbiont from *H. acanthopus* was aligned to the 39 contigs of the *Neisseria*-related symbiont from *P. serrata*, yielding 27 Locally Collinear Blocks (Figure 1). The resulting synteny regions are explored in more detail in Supplementary Data 1. To provide a comparison of selected metabolic pathways, we extended the analysis from our previous work on *Legionella polyplacis* (Rihova et al 2017) which followed the selection of genes published by (Moran et al 2008). The comparison was based on the RAST annotation for the two *Neisseria*-related symbionts, and the KEGG database (Kanehisa et al 2016) for our other bacteria.

**Figure 1.**
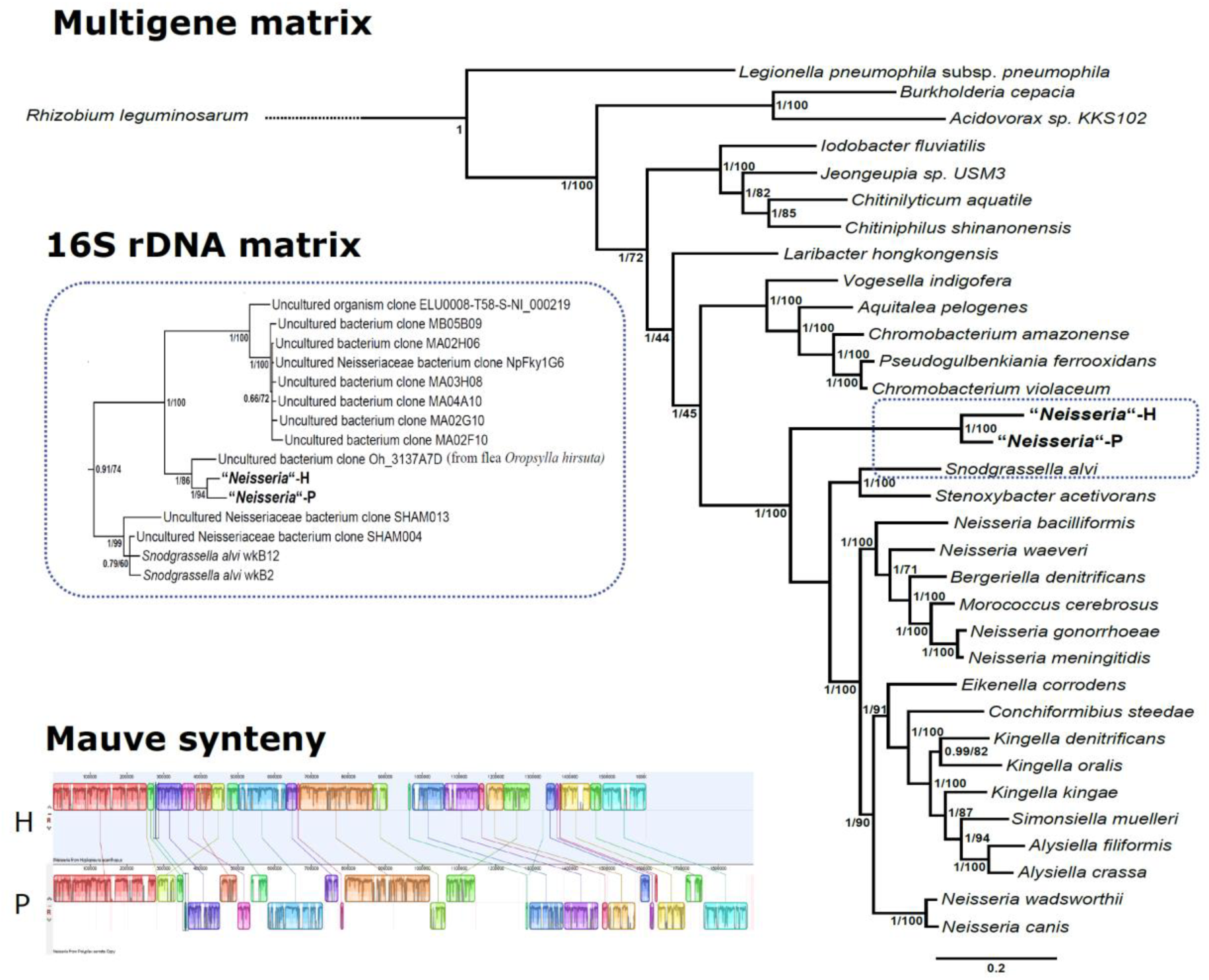
Phylogenetic relationships of the two *Neisseria*-related symbionts. Multigene matrix: Bayesian analysis of the multigene matrix; the numbers at the nodes show posterior probabilities/bootstrap supports obtained by the Maximum Likelihood analysis in PhyML. 16S matrix: part of the tree obtained by the Bayesian analysis of the 16S matrix showing relationships of the two *Neisseria*-related symbionts to several Uncultured bacteria (see Supplementary Figure1 for the complete tree); the numbers at the nodes show posterior probabilities/bootstrap supports obtained by the Maximum Likelihood analysis in PhyML. Mauve synteny: an overview of strong synteny between the two symbionts; H - “*Neisseria*”-H, P - “*Neisseria*”-P (see Supplementary Data 1 for a complete list of the genes).

### Amplicon library preparation and sequencing

Samples of 190 *Polyplax serrata*, 14 *Hoplopleura acanthopus*, and 2 *Hoplopleura edentula* were collected from different rodent species and populations from 2014-2018. Details on sample origin, including the sampling locations, GPS coordinates, and rodent host species, are provided in Supplementary Data 2. DNA templates were extracted from each individual using QIAamp DNA Micro Kit (Qiagen). Two independent multiplexed 16S rRNA gene amplicon libraries were built for each of the louse genera. *Polyplax serrata* gDNA from 190 samples was amplified using fused barcoded primers of the EMP protocol (http://www.earthmicrobiome.org/protocols-and-standards/16s/) producing approximately 300-350 bp long amplicons of the V4 hypervariable region. Complete barcode assignation is available in Supplementary Data 2. The pooled library included 16 negative controls for the extraction procedure and PCR amplification. We sequenced the library in a single MiSeq run using v2 chemistry with 2 × 150 bp output. Amplicons exceeding 450bp of the 16S rRNA gene were retrieved for *Hoplopleura* species using a double barcoding strategy with 515F and 926R primers (Parada et al 2016, Walters et al 2016) barcodes and primers available in Supplementary Data 2. Sixteen *Hoplopleura* samples were part of a pooled library of 432 samples containing 8 negative extraction and amplification controls as well as 6 positive controls. The positive controls comprised 3 samples of commercially available mock communities with equal composition of 10 bacterial species and 3 samples with a staggered profile (ATCC® Microbiome Standards). The data were generated in a single MiSeq run using v3 chemistry with 2 × 300 bp output.

### Processing and analyses of 16S rRNA gene amplicons

The amplicons, originating from two different library designs, were processed as two independent data sets. The raw data were quality checked in FastQC and trimmed (needed for reverse reads from 300bp output mode) using USEARCH v9.0.1001 (Edgar 2013). The reads were further processed according to the following workflow implementing suitable scripts from USEARCH v9.0.1001. Pair-end reads were demultiplexed (-fastx_demux), merged (-fastq_mergepairs), and trimmed (-fastx_truncate) in order to remove the primer sequences, then quality filtered (-fastq_filter). The final length of *Polyplax* and *Hoplopleura* amplicons were 253 bp and 410bp, respectively. The two datasets were clustered at 100% identity providing a representative set of sequences for *de novo* OTU picking. OTUs were defined using the USEARCH global alignment option at 97% identity. We used the blast algorithm (Camacho et al 2009) against SILVA 132 database (Quast et al 2013) to assign taxonomy of the representative sequences. Chloroplast, mitochondrial OTUs, OTUs of extremely low abundance (as recommended by Bokulich et al 2013), and singletons were filtered out from the final OTU table using QIIME 1.9 (Caporaso et al 2010).

Since the negative controls for the *Polyplax* dataset contained a considerable number of bacterial reads, 9299 on average compared to 161 in *Hoplopleura* library, the OTU tables were filtered for potential contaminants. These were defined as OTUs comprising more than 1% of reads in any negative control found in more than one fourth of negative controls for both *Polyplax* and *Hoplopleura* libraries. Two such OTUs were filtered out from the *Polyplax* dataset (OTU_2 assigned as *Staphylococcus* sp. and OTU_4 assigned as *Acinetobacter* sp). For the *Hoplopleura* dataset we discarded three OTUs. Two were found in the negative controls (OTU_33; *Geobacillus* sp. and OTU_1; *Pectobacterium* sp.) and one (OTU_4; *Staphylococcus* sp.) originated in the mock communities. Since the presented analysis centers on symbiotic taxa in the microbiome, only highly abundant taxa were considered, i.e. OTUs comprising ten or more percent of the reads within a particular sample. Under the assumption of high symbiont prevalence within the host populations, our final selection of taxa includes OTUs that occur in more than 5 individuals across the analyzed *Polyplax* samples, or in at least two *Hoplopleura* samples.

### Host phylogenetic background

We used 379 long sequences of the COI gene (amplified with L6625 and H7005 (Hafner et al 1994) primers) to determine phylogenetic background of 190 *Polyplax serrata* samples. For phylogenetic reconstruction of *Hoplopleura* host species we amplified 968 bp long region of the COI gene using LCO1490 (Folmer et al. 1994) and H7005 primers. PCR products were enzymatically purified and sent for Sanger sequencing. All sequences are available in GenBank under accession numbers provided in Supplementary Data 2. The matrices were aligned using E-INS-i algorithm of MAFFT v7.450 (Katoh et al 2002) in Geneious software. Ambiguously aligned positions and divergent blocks were discarded using Gblocks v. 091b (Castresana 2000). Phylogenetic relationships were reconstructed by Bayesian inference with the GTR +I+G best-fit model selected according to a corrected Akaike information criterion using jModelTest2 v2.1.10 (Darriba et al 2012, Guindon and Gascuel 2003). *Polyplax spinulosa* was used as the outgroup for the *Polyplax serrata* dataset and the *Polyplax serrata* sequence was used as the outgroup for *Hoplopleura* spp. dataset. Bayesian analyses conducted in MrBayes v3.2.4 (Ronquist et al 2012) consisted of two parallel Markov chain Monte Carlo simulations with four chains run for 10 million generations and sampling frequency of 1,000 generations. The convergence of parameter estimates and their ESS values was checked in software Tracer v1.7 (Rambaut et al 2018). Two and a half million generations (25%) were discarded as burn-in.

### Phylogenetic origin of the symbionts

Phylogenetic analysis of the *Neisseria*-related symbionts was performed on two different matrices, the concatenated “multigene matrix” and the “16S matrix”. To avoid a possible artefact due to HGT, the “multigene matrix” was composed of 10 genes with reliably supported origin within Neisseriales (the genes selected arbitrarily from the blast category “Neisseriales”; see above) which were present in both genomes. Two betaproteobacteria of the order Burkholderiales, *Burkholderia cepacia* and *Acidovorax* sp. KKS102, one gammaproteobacterium, *Legionella pneumophila subsp. pneumophila str. Philadelphia 1*, and one alphaproteobacterium, *Rhizobium leguminosarum*, were used as outgroups (Supplementary Data 3). For each gene, the sequences were aligned in MAFFT v7.450 using the E-INS-i setting. Ambiguously aligned positions and divergent blocks were discarded using Gblocks v. 091b. The LG +G+I was determined as best fitting model for all matrices by Akaike information criterion (AIC) using smart model selection of PhyML (Lefort et al 2017). Maximum-likelihood phylogenetic reconstructions were performed using online PhyML server v3.0 (Guindon et al 2010) with 100 bootstrap replicates for each single-gene alignment and also for the concatenated “multigene matrix”. Bayesian inference of the “multigene matrix” was conducted in MrBayes v3.2.5 using LG +G+I evolutionary model. Four chains were run for 20,000,000 generations with sampling frequency set to 1,000 generations. Convergence was checked in Tracer v1.7.

The “16S matrix” was designed with the aim to obtain wider phylogenetic context by including the bacteria for which the 16S rRNA gene sequence is the only available marker. The 16S rRNA gene sequences were retrieved by blastn from the GenBank (Supplementary Data 3). Two betaproteobacteria *Taylorella equigenitalis* str. 09-09 and *Advenella kashmirensis* str. cv4, and one alphaproteobacterium, *Rhizobium capsici* str. IMCC34666, were used as outgroups. The matrix was prepared with the same procedure as our “multigene matrix” and analyzed by maximum likelihood (ML) and Bayesian inference (BI). The evolutionary models best fitting to dataset were selected according to the Akaike information criterion (AIC) using jModelTest2 v. 2.1.10. ML analysis and 100 bootstrap replicates were performed with online PhyML using selected TN93 +G+I evolutionary model. BI analysis was performed in MrBayes v. 3.2.5, using GTR +G+I substitution model running four chains for 10,000,000 generations and checked for convergence as was previously described.

### Fluorescence in situ hybridization (FISH) and microscopy

We investigated the spatial distribution of the symbionts within the body of *Hoplopleura acanthopus* using whole-mount FISH. First, we fixed tissues by incubating at least three specimens of the louse in 4% paraformaldehyde solution at 4°C for 39 h. Subsequently, insects were transferred and kept at 4°C in Carnoy’s solution for 27 h, 2% hydrogen peroxide ethanol solution for 3 days and 6% hydrogen peroxide ethanol solution for 10 days to quench tissue autofluorescence. We then washed specimens with 400 µl of hybridization buffer (900 mM NaCl, 20 mM Tris-HCl pH 7.4, 0.01% sodium dodecyl sulfate, SDS, and 30% formamide) at 46°C for 10 min, pre-hybridized with 200 µl of hybridization buffer at 46°C for 1 h, and hybridized at 46°C for 3 h with 400 µl of hybridization buffer containing probes. All fluorescent probes were obtained from Sigma-Aldrich (Germany) and used at a concentration of 2.5 ng μl^−1^ for Cyanine 5 (Cy5)-labelled EUB338 (5’-GCTGCCTCCCGTAGGAGT - 3’, targeting all bacteria (Amann et al 1990) and 6-carboxyfluorescein (6-Fam)-labelled beta-572 (5’-TTAACCGTCTGCGCTCGCTT −3’, targeting the family Neisseriaceae (Martinson et al 2012). We pre-evaluated the required stringency of the hybridization conditions in silico using mathFISH (Yilmaz et al 2011). Following hybridization, we washed specimens twice with 400 µl of pre-warmed wash buffer (20 mM Tris-HCl, 5 mM EDTA, 0.01% SDS and 112 mM NaCl) at 48°C for 10 min. All incubations were carried out with ongoing shaking at 300 rpm. We then placed lice on microscope slides, incubated them in ~50 μl of 4’,6-diamidino-2-phenylindole (DAPI) solution (1 ng μl-1) in the dark for 10 min, and mounted slides in Mowiol anti-fading medium (Kuraray Europe GmbH, Japan). We captured the fluorescent signals with a laser scanning confocal microscope Olympus FV3000 (Olympus, Japan). We acquired at least three confocal stacks (up to 30 scans per optical slice) at 100x, 400x and 630x magnifications, a color depth of 24 bit and a resolution from 1 to 2 μm per pixel (depending on the fluorochrome) by investigating multiple regions from each of the three replicate specimens. Resultant images were processed with the ImageJ distribution Fiji (Schindelin et al 2012, Schneider et al 2012).

## Results

### Genomes of the Neisseria-related symbionts

The complete closed genome of the obligate *Hoplopleura acanthopus* symbiont, obtained with a combination of Nanopore and Illumina data, is 1,607,498 bp long with 33.4% GC content. It contains 1421 protein encoding genes, 9 genes coding rRNAs, and 39 genes coding tRNAs (Supplementary Data 1). The 9 rRNA genes represent 3 complete 16S-23S-5S rRNA gene operons, but they are arranged in an unlinked manner known from many other bacteria (Brewer et al 2019), including other P-symbionts (Munson et al 1993). The 23S rRNA and 5 rRNA genes are placed in close proximity, while the 16S rRNA gene is separated by long stretches of DNA. The genome draft of *Polyplax serrata* symbiont consisted of 39 contigs which sum to 1,839,042 bp, with the GC content 34.0%. In this assembly, we identified 1789 protein encoding genes and 37 tRNA genes (Supplementary Data 1). Due to the fragmentation of the genome in contigs, the rRNA genes were not reliably assembled and their number remains unclear. To make the following text intelligible, we recategorise Neisseria-related symbionts with the terms *“Neisseria”-H* (symbiont of *Hoploplera*) and *“Neisseria”-P* (symbiont of *Polyplax*). Genome size and content of these two symbionts, in comparison to other lice symbionts (*Riesia* spp., *Puchtella pedicinophila*, *Legionella polyplacis*, and the symbiont from *Proechinophthirus*), *Snodgrassella* and *Neisseria*, are summarized in Table 1. When blasted against the NCBI nr database, a large proportion of the genes in both *Neisseria*-related symbionts did not yield Neisseriales within the best hits (see Methods and Supplementary Data 1). This is in remarkable contrast to *Legionella polyplacis*, for which 96.3% of the genes were assigned to *Legionella* by the blast search. However, when compared to each other, the two *Neisseria*-related symbionts share a high proportion of their genes and even display a considerable degree of synteny (Supplementary Data 1). When aligned with Mauve software the syntenies were placed into 27 LCBs, the longest blocks extending 100 kb and hundred genes (Figure 1).

#### Phylogeny

In both phylogenies (the 16S rRNA gene and the multigene) the two *Neisseria*-related symbionts cluster as sister taxa on a long common branch, placed firmly within Neisseriales with high nodal support. In the more robust multigene analysis the pair branches as an isolated offshoot at the base of Neisseriaceae (Figure 1). The analysis of 16S rRNA gene, for which a broader taxonomic spectrum is available, revealed additional close relatives which could not be included into the “multigene matrix” due to lack of data (Supplementary Figure 1; detail in Figure 1). All of them have been described as “*Uncultured bacterium*” from human and insect samples (Oteo et al 2014). In contrast to the “multigene matrix”, the “16S matrix” placed the pair of *Neisseria*-related symbionts into a monophyletic cluster together with *Snodgrassella*.

#### Amplicon based screening

Sequencing of multiplexed 16S rRNA gene libraries produced high quality amplicon data. Similar numbers of merged 16S rRNA bacterial sequences were retrieved across individual *Polyplax* and *Hoplopleura* samples (21, 205 and 20, 201 on average). For the even mock communities included in the *Hoplopleura* library as positive controls (see Methods), the sequencing recovered all 10 bacterial taxa in comparable abundances. The data from the staggered communities, designed for testing sequencing sensitivity and PCR bias, confirmed that our approach can reveal complete microbiome profiles, including low abundant taxa. In particular, sequences of *Bifidobacterium adolescentis (ATCC15703)* and *Deinococcus radiodurans (ATCC BAA-816)*, both present in 0.04% of the original mock DNA template, comprised on average 0.02% and 0.01% of the reads among the three sequenced mock communities. However, the staggered composition of the three mock samples did result in a preferential amplification of one of two dominating taxa, i.e. *Staphylococcus epidermidis* (ATCC 12228). Compared to the mock template where this taxon comprises 44.78% of the total DNA, the average read abundance equaled 70.12%. Therefore, no quantitative analyses of 16S rRNA gene amplicons were used in this study.

#### Distribution of dominant bacterial taxa

Microbiomes of the tested species/lineages of the lice are dominated by several bacterial taxa with a complex distribution pattern. For two of these bacteria, *Legionella polyplacis* from *Polyplax* and Neisseriaceae from *Hoplopleura* lice, their symbiotic nature could be clearly demonstrated based on the complete genome characteristics and distribution within the host. For other bacteria, partial 16S rRNA gene amplicons provide two (not entirely independent) kinds of information: taxonomical assignment and GC content. Besides Neisseriaceae, two more OTUs, identified as dominant taxa (see methods for definition) and taxonomically assigned to *Blochmania* and *Arsenophonus* genera, were associated with *Hoplopleura* samples (Figure 2). In addition to the sequence characteristic (GC content of 43.9% and 49.3%), the distribution of these OTUs among the two *Hoplopleura* species and *H. acanthopus* populations points to their symbiotic nature.

**Figure 2.**
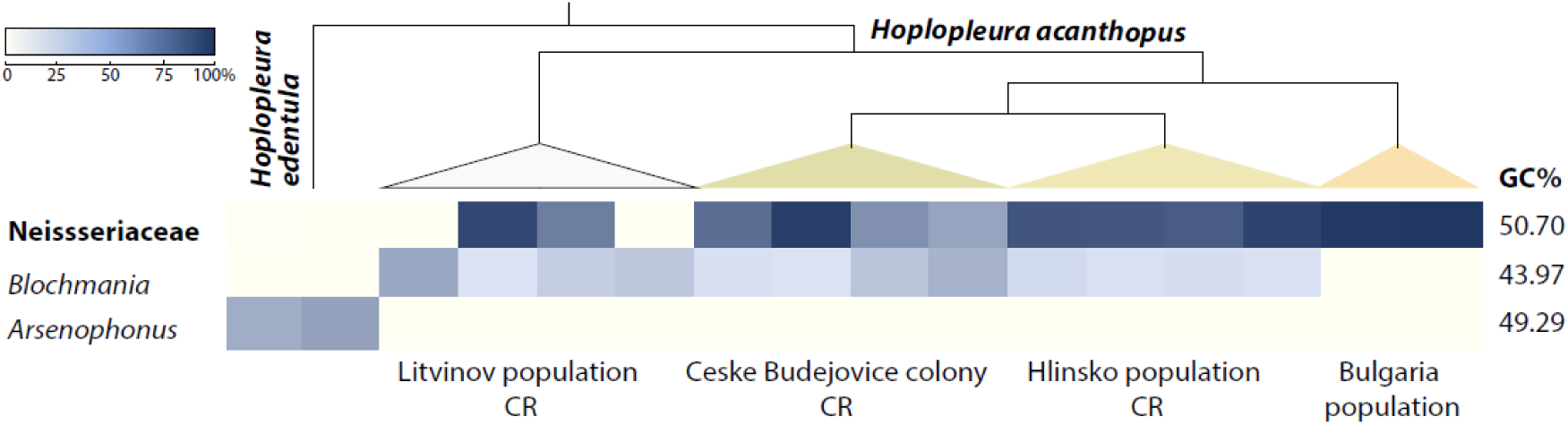
Highly abundant taxa found in *H. edentula* and four different populations of *H. acanthopus* using high throughput 16S rRNA gene amplicon sequencing. Detailed phylogenetic relationships of the hosts are shown in the Supplementary Figure 2.

For *Polyplax serrata* a comprehensive population-wide amplicon screening revealed (besides *L. polyplacis*) 9 dominant taxa assigned to the genus or family level (Figure 3). The distribution of OTUs with a low GC content, i.e. *Buchnera* (45.1% GC) and *Arsenophonus* 2 (49.4% GC), reflects the genealogy of the host and thus indicates putative obligate coevolving symbionts. For the other taxa, the taxonomic assignment and GC content >50% (with the exception of *Cloacibacterium*) indicates that the bacteria may represent environmental contamination or very early symbiotic associations, e.g. Neisseriaceae taxon and *Arsenophonus* 1. However, it is important to note that the taxonomical assignment of the OTUs are based on a short sequence and should be interpreted as approximate affiliations rather than precise phylogenetic position, particularly compared to highly derived genomes like *Buchnera* and *Blochmannia*.

**Figure 3.**
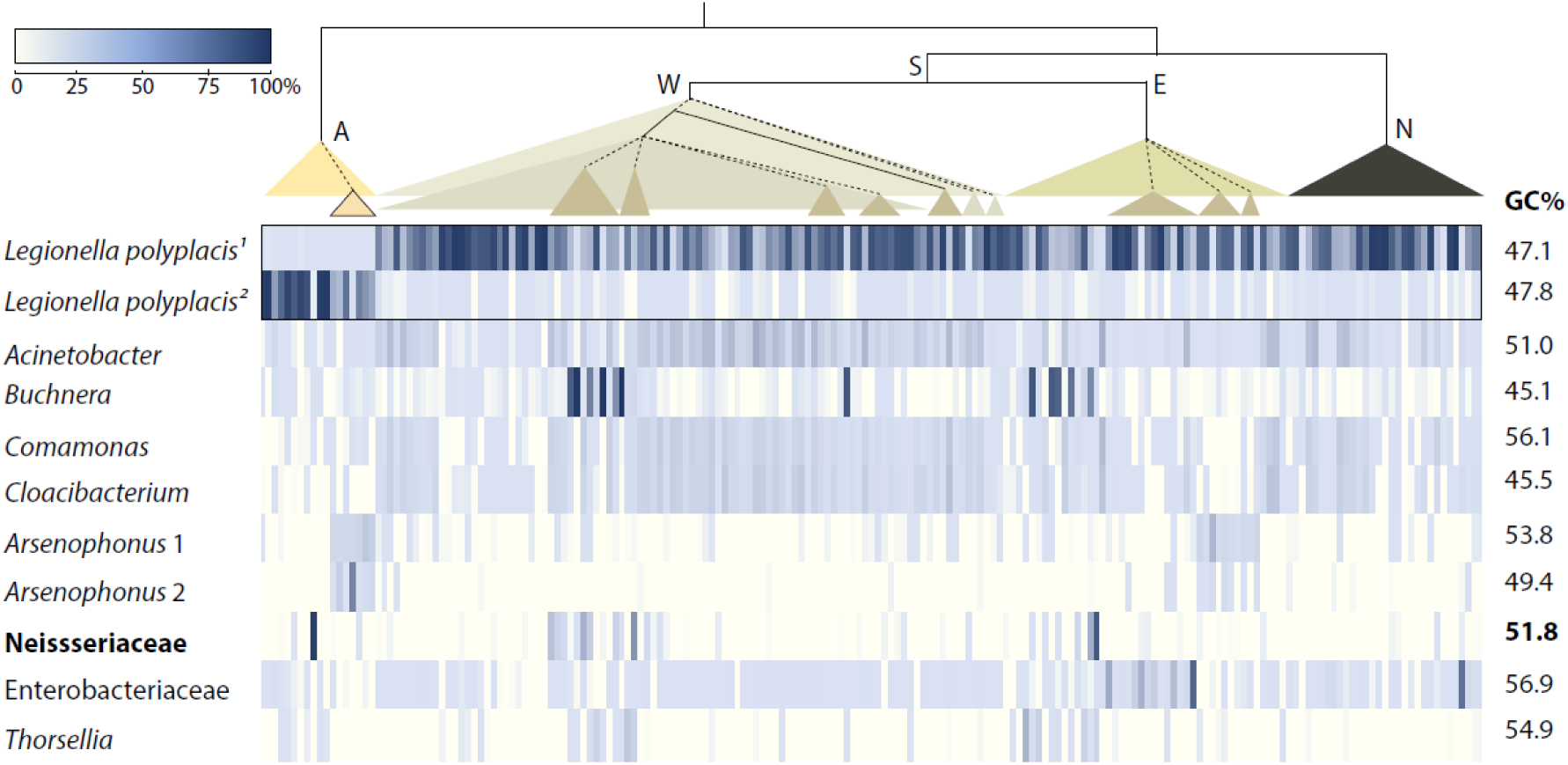
Highly abundant taxa found in *Polyplax serrata* samples from distinct populations using high throughput 16S rRNA gene amplicon sequencing. The phylogenetic scheme simplifies the COI based phylogeny of individual samples provided in Supplementary Figure 3. Designation of the branches is based on mtDNA structure described in the study by (Martinu et al 2018): A = lineage specific to *Apodemus agrarius*, S = lineage specific to *Apodemus flavicollis* (W = west sublineage, E = east sublinage), N= nonspecific lineage from *A. flavicollis* and *A. sylvaticus*. The numbers for *Legionella polyplacis* designate two different OTUs, reflecting evolutionary changes accumulated after the split of *Polyplax serrata* lineages.

#### Localization of the lice symbionts

Both the specific Neisseriaceae and the universal bacterial probes hybridized to bacteria located within DAPI-stained cells (~20-30 μm). In females, these putative bacteriocytes formed weakly adherent clusters found above ovarial ampullae. The two probes did not provide entirely overlapping patterns (Figure 4). However, it is difficult to decide whether this difference is due to presence of two different bacteria within the bacteriocytes or due to the difference in signal intensity produced by chemically distinct chromophores. Many bacteriocytes containing intracellular bacteria were also found in the posterior part of abdomen in male lice (Supplementary Figure 4).

**Figure 4.**
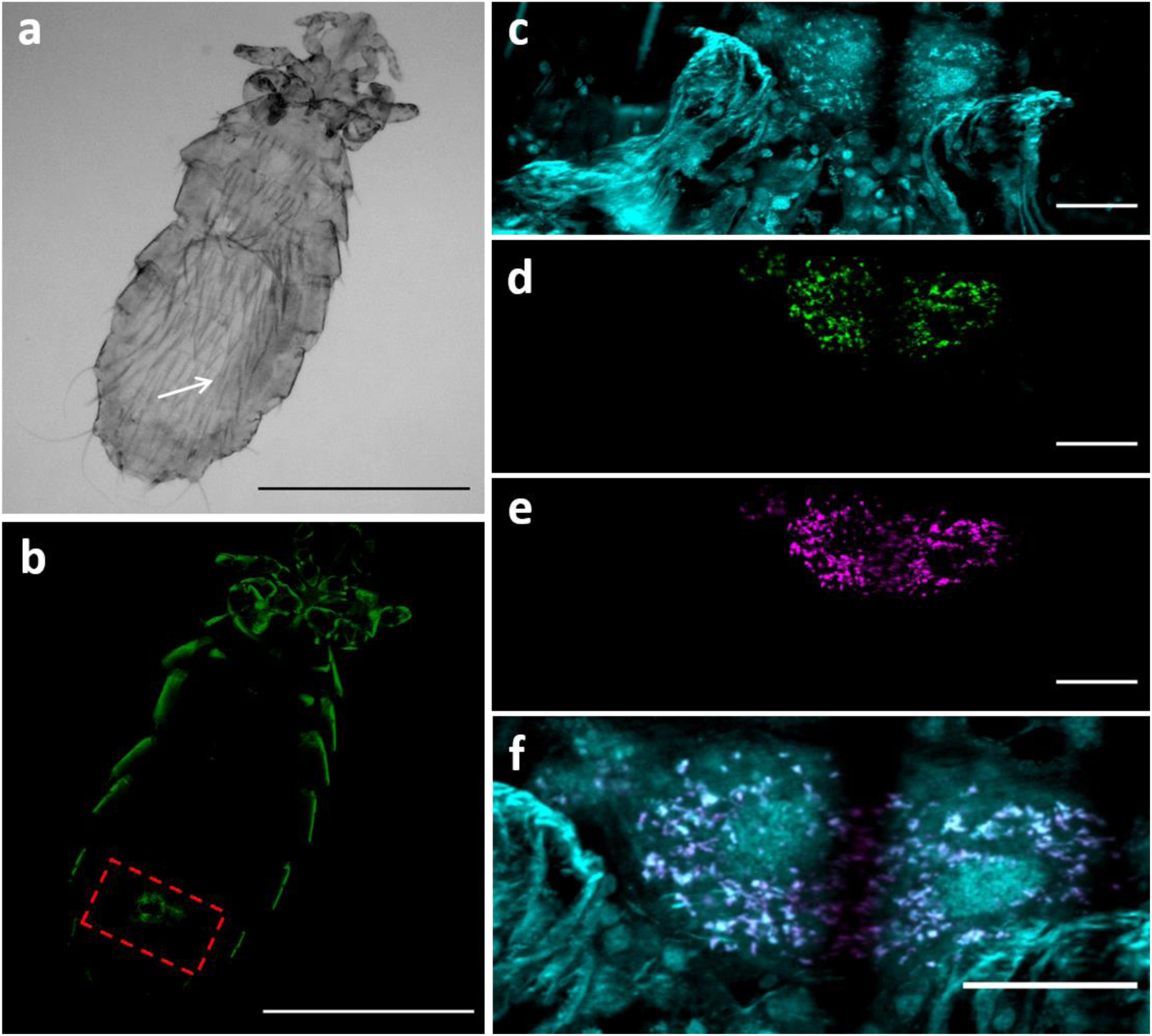
Light and confocal microscopy of a FISH-stained female *H. acanthopus*. (a) Light microscopy image showing the louse body containing a developing egg (white arrow). (b) Hybridization signal for the Neisseriaceae specific probe beta-572 (green) shows the localization of the symbionts. The dashed red square defines the region shown in panel c, d and e. (c, d and e) Hybridization signals for DAPI (cyan), the Neisseriaceae specific probe beta-572 (green) and the generic bacterial probe EUB338 (magenta) are shown in panel c, d and e, respectively, and were combined in the merged color image (f), with the *Neisseria*-related symbionts resulting being in white. DAPI staining in panel c defines also the localization of the bacteriocytes above ovarial ampullae. Scale bars: (a, b) 500 μm, (c, d, e and f) 20 μm.

## Discussion

The results of the combined genomic and amplicon analyses which we present here, illustrate a positive relationship between the dynamics of louse microbiomes and the observed high diversity of symbiotic bacteria in lice (Allen et al 2016, Boyd et al 2016, Boyd et al 2014, Fukatsu et al 2009, Hypsa and Krizek 2007). The two *Neisseria*-related symbionts, which are the primary focus of this study, represent a novel lineage, extending the known phylogenetic span of louse symbionts. Considering their genomic characteristics together with their distribution across the lice populations, we hypothesize that they are bacteria in an early/intermediate stage of evolution towards obligate symbiosis, which established their symbiosis independently within two different genera of lice. Consistent with this view, FISH analysis highlighted their localization within putative bacteriocytes (Figure 4 & Supplementary Figure 4), and amplicon screening showed that they are part of a rich microbiomes diversity (Figures 2 & 3), in which some of the other dominant taxa are closely related to the other known obligate symbionts in insects (the most degenerated ones may even be disappearing ancient P-symbionts). This complex picture suggests dynamic turnover of the symbiotic bacteria, their frequent acquisitions, losses and replacements even within local louse populations and young phylogenetic lineages.

### Genomic characteristics of the *“Neisseria”* symbionts suggest an intermediate stage of the evolution

We propose that the *Neisseria*-related symbionts are in an early or intermediate state of symbiosis evolution. We base this view on comparisons with other known symbionts. Full genomes are currently available for four lineages of lice-associated symbionts. Three are highly reduced and display strong compositional shift towards AT, typical for many P-symbionts (Table 1). The fourth possesses a large genome and is supposedly a recent acquisition (Boyd et al 2016). The genomes of the two *Neisseria*-related symbionts can thus be compared to various lice P-symbionts and to “free living” members of Neisseriales. Both “*Neisseria*”-H and “*Neisseria*”-P display significantly weaker genome degeneration than the highly reduced *Riesia*, *Puchtella*, and *Legionella* (Boyd et al 2017, Rihova et al 2017). They have higher GC content, considerably larger genomes (approximately three times higher number of genes), and consequently more complete metabolic pathways (Supplementary Data 1). On the other hand, their genomes are recognizably reduced and the GC content decreased when compared to their relatives (e.g. the genus *Neisseria*, but also the bee symbiont *Snodgrassella*) or to the presumably young *Sodalis*-like symbiont from the louse *Proechinophthirus fluctus* (Boyd et al 2016). The comparison of selected metabolic pathways shows that the *Neisseria*-related symbionts retain considerably greater numbers of genes in various categories than the reduced genomes of *L. polyplacis* and *R. pediculicola* (Supplementary Data 1). This difference is particularly strong in the *Recombination and repair* category (as one example) but also in several amino acid biosynthesis pathways. An interesting example of differences between the two *Neisseria*-related symbionts is the complete histidine pathway in the “*Neisseria-*P” symbiont, entirely lost in the more reduced “*Neisseria-*H” (as well as in *L. polyplacis* and *R. pediculicola*). Another conspicuous difference between the *Neisseria*-related symbionts and the other two louse symbionts regards their capacity to build the cell walls. Similar to the *L. polyplacis* and *R. pediculicola*, the *Neisseria*-related symbionts retain the path for peptidoglycan synthesis, but unlike them, they also possess the genes for penicillin binding protein class A and a complete pathway for lipid A, required for synthesis of lipopolysaccharide. In contrast, both *Neisseria*-related symbionts seem to lack the rod-shape coding genes. Finally, the genomes of both *Neisseria*-related symbionts retain various genes connected to DNA exchange and/or transport, such as mobile elements, type iv pili, and secretion systems (Supplementary Data 1). If, as suggested by many of the described genomic features, “*Neisseria”*-H is a young symbiont in an early state of evolution, we could assume that it only recently replaced a more ancient P-symbiont, which was fulfilling the nutritional role prior to the acquisition of the *Neisseria*-related symbiont. As discussed below, such a putative P-symbiont was indeed detected in the majority of the *H. acanthopus* microbiomes. Also, the FISH survey indicated that the bacteriocytes might be inhabited by two different bacteria. However, this interpretation should be taken with caution since the non-overlapping signal may reflect different properties of the used chromophores (Figure 4).

### Distribution and origin of the symbionts in the lice microbiomes

Since the process of genome degradation starts once the bacterium becomes an obligate vertically-transmitted symbiont we should expect, at least during the initial phase of the symbiogenesis, a correlation between the degree of genome degradation and the duration of host-symbiont coevolution (Moran 1996). For example, among the symbionts of sucking lice, the high degree of genome degradation of *R. pediculicola* and *L. polyplacis* indicates a relatively long and intimate association with the host. In correspondence with this presumption, the *Riesia* lineage has been found in several louse species of two different genera, *Pediculus* and *Phthirus*, and *L. polyplacis* is hosted at least by two louse species, *P. serrata* and *P. spinulosa* (Hypsa and Krizek 2007). Moreover, our extensive amplicon screening shows that *L. polyplacis* is consistently present in a broad geographic and phylogenetic sample of *P. serrata* as a dominant bacterium (Figure 3). When compared to these two well documented examples of established P-symbionts, the *Neisseria*-related symbionts, with an intermediate degree of the genome degeneration, show more restricted and patchy distribution (Figure 3). Only in *H. acanthopus* were they consistently present (the Neisseriaceae OTU) as the most dominant bacterium (with the exception of two specimens from one population; Figure 2), but were not found in the two examined specimens of the related species (*H. edentula)*. The overall variability of the microbiomes was apparently correlated with the lice genetic background: in Bulgarian samples the Neisseriaceae OTU was the only present bacterium, in other populations of *H. acanthopus* it was accompanied by an unknown bacterium which the blast search affiliated with *Blochmania*, and in the two samples of *H. edentula* the only present OTU corresponded to the genus *Arsenophonus.* Since *P. serrata* is known to harbor the typical obligate P-symbiont *Legionella polyplacis* (Rihova et al 2017), we screened this louse more extensively across several populations and genetic lineages. The results confirmed a ubiquitous presence of *L. polyplacis*, which in most cases was the most abundant OTU, and only occasional co-occurrence of the Neisseriaceae OTU (Figure 3). The split of *L. polyplacis* into two different OTUs correlated to the host’s phylogeny, reflects evolutionary changes during the evolution of the symbiont in distant host lineages, but certainly does not suggest the presence of two independent symbiotic lineages (in fact phylogenetic and genomic analyses confirm that *Polyplax*-*Legionella* co-evolution crosses the host species boundaries and the same symbiont is also present in the related louse species *P. spinulosa;* (Hypsa and Krizek 2007). Three additional OTUs in *P. serrata* microbiomes show affinity to known insect symbionts, the *Buchnera* OTU and two *Arsenophonus* OTUs. Based on their genetic divergence and the differences in their GC content, the two *Arsenophonus* OTUs seem to represent two different lineages.

In respect to the general concept of symbiont acquisition, loss, and replacement within insects, and the high dynamism of louse microbiomes, two OTUs are of particular interest. Both the *Buchnera* OTU from *P. serrata* and the *Blochmania* OTU from *H. acanthopus* seem to represent strongly derived symbiotic genomes (Figure 2 & 3), which blast-assigned taxonomy reflects the low GC content rather than real phylogenetic relationships. Since our metagenomic data did not yield any reliable information on either of these bacteria, it is difficult to hypothesize about their phylogenetic origin and function in the host. However, strong genome reduction, deduced from the GC content of the 16S rRNA gene amplicon, suggests that they may represent the scattered remains of ancient symbionts, now retreating from the host’s population and replaced with more recent acquisitions. Interestingly, the FISH analysis shows that apart from the “*Neisseria*-H” symbiont, the bacteriocytes of *H. acanthopus* harbor another bacterium (Figure 4). Since the metagenomic assembly did not contain any other bacterial contigs, we were not able to identify the origin of this second symbiotic bacterium.

Considering this distribution pattern and the low degree of genome modifications, it is unlikely that occurrence of the two “*Neisseria* symbionts” in the two different lice lineages is due to a common symbiotic origin in the *Hoplopleura*-*Polyplax* ancestor. The most parsimonious explanation is thus an independent origin of the symbiosis in each louse genus. The close phylogenetic relationships of the two symbionts (sister taxa in the phylogenetic tree) poses an interesting question on the underlying mechanisms. Co-occurrences of closely related symbiotic bacteria in related insect hosts are usually consequences of either co-speciation or a tendency of specific bacterial lineages to frequently establish symbiosis with specific insect hosts (e.g. *Arsenophonus*, *Wolbachia*). However, neither of these explanations can be applied to the lice-*Neisseria* association. Members of the family Neisseriaceae are only rarely found in symbiotic association with insects. The only well documented case of obligate symbiosis is the genus *Snodgrassella* found in several species of bees and bumblebees (Kwong and Moran 2013). Based on the 16S rRNA gene phylogeny, the closest relative of the louse-associated *Neisseria* is an uncultured bacterium described from a flea *Oropsylla hirsuta* (Jones et al 2008), for which no other information is currently available. The two *Neisseria*-related symbionts are not only a novel lineage within the diversity of lice-associated bacteria, they are also the first example of louse symbionts originating outside Gammaproteobacteria. Identification of the main factors behind the repeated establishment of symbiosis between a louse and the *Neisseria*-related bacterium is difficult with the available information. It is interesting to note that, similar to *Legionella polyplacis*, the *Neisseria*-like symbionts originate from a bacterial lineage which is rarely found in insects and is a well-known vertebrate pathogen. Phylogenetic correlation between the vertebrate pathogens and symbionts of blood-feeding arthropods was previously reported in ticks (Ahantarig et al 2013, Felsheim et al 2009, Guizzo et al 2017, Niebylski et al 1997, Noda et al 1997). For one of the tick symbionts, *Francisella*-like bacterium, the origin from mammalian pathogen was recently suggested by (Gerhart et al 2016).

## Acknowledgments

This work was supported by the Grant Agency of the Czech Republic (grant GA18-07711S to V.H.). We would like to acknowledge the sequencing services of Genomics Core Facility, EMBL Heidelberg, Heidelberg, Germany and the DNA Services of W.M Keck Center, University of Illinois at Urbana-Champaign, IL, USA. Access to computing and storage facilities owned by parties and projects contributing to the National Grid Infrastructure MetaCentrum provided under the programme “Projects of Large Research, Development, and Innovations Infrastructures” (CESNET LM2015042), is greatly appreciated. In addition, we thank Joel J. Brown for his language corrections on this manuscript.

## Supplementary Information

***Supplementary Data 1:*** Complete genomes of the *Neisseria*-related symbionts with highlighted synteny; summarization of blast categories; overview of biosynthetic pathways (available at https://www.dropbox.com/sh/frkzmjwnnzrc3v3/AAAWomM8m_8LIMM2KK57cXH-a?dl=0)

***Supplementary Data 2:*** Complete sample metadata including accession numbers for host markers. (available at https://www.dropbox.com/sh/frkzmjwnnzrc3v3/AAAWomM8m_8LIMM2KK57cXH-a?dl=0)

***Supplementary Data 3:*** Accession numbers of the sequences used for bacterial phylogenies. (available at https://www.dropbox.com/sh/frkzmjwnnzrc3v3/AAAWomM8m_8LIMM2KK57cXH-a?dl=0)

***Supplementary Figure 1:*** Consensus of phylogenetic tree derived from the “16S matrix”.

***Supplementary Figure 2:*** Phylogeny of *Hoplopleura* lice.

***Supplementary Figure 3:*** Phylogeny of *Polyplax serrata* lice.

***Supplementary Figure 4:*** Confocal microscopy of a FISH-stained male *H. acanthopus*.

---

**Supplementary Figure 1:**
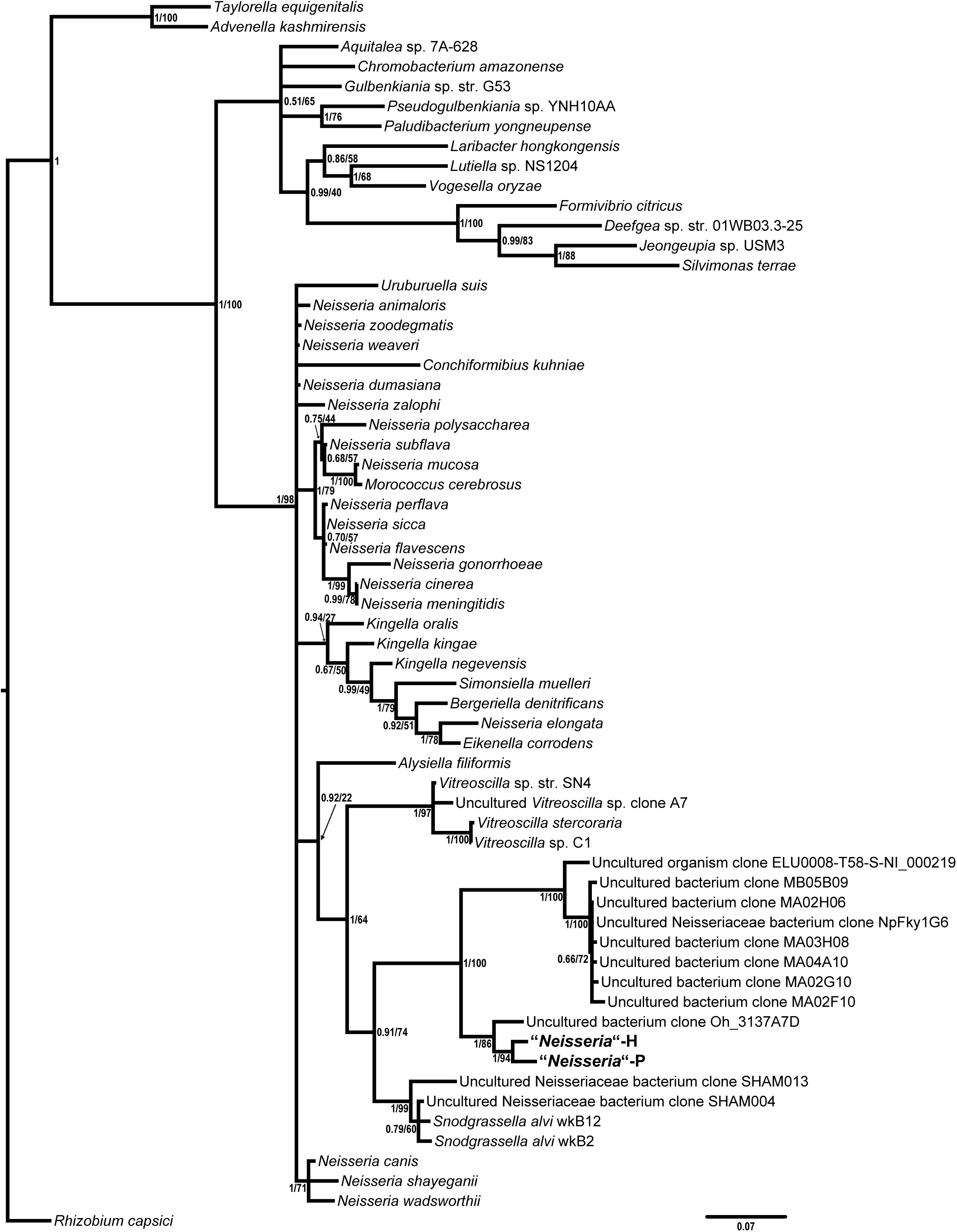
Consensus of phylogenetic trees derived from the “16S matrix” by Bayesian inference (MrBayes) and Maximum Likelihood (PhyML). The values at the nodes show statistical supports (Bayesian posterior probability/maximum likelihood bootstrap).

**Supplementary Figure 2:**
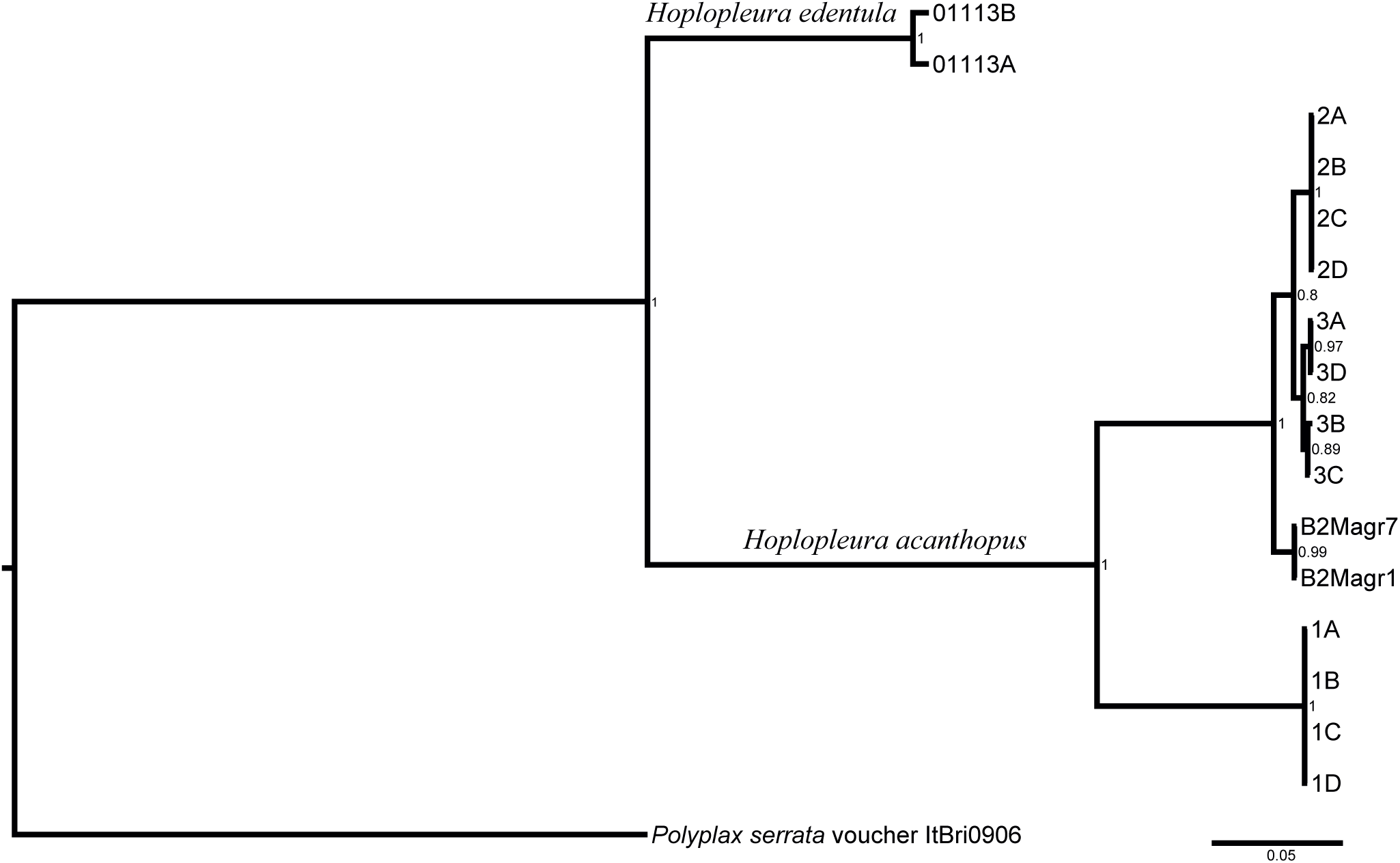
Phylogeny of *Hoplopleura* lice obtained with Bayesian inference (MrBayes). The values at the nodes show statistical supports (Bayesian posterior probability).

**Supplementary Figure 3:**
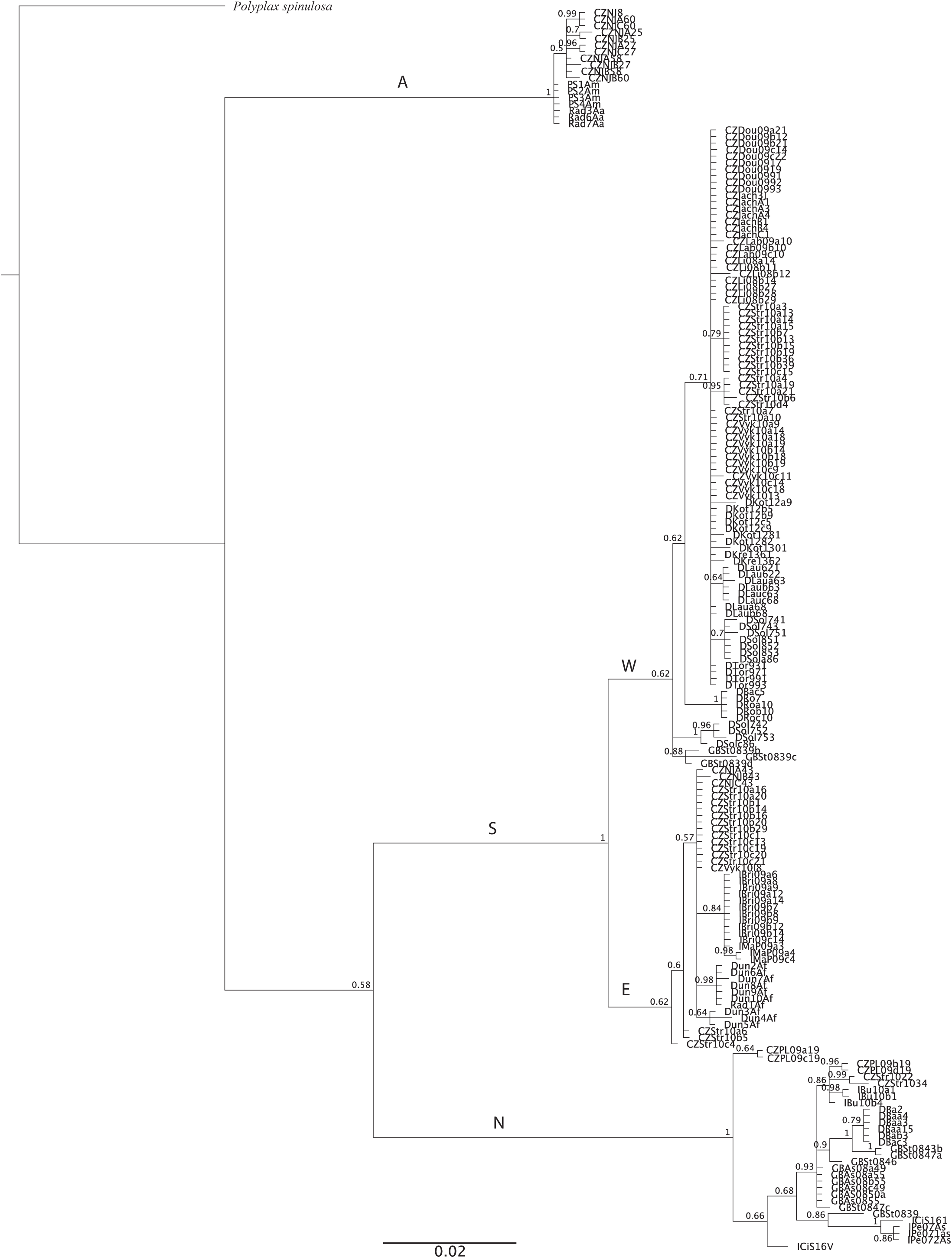
Phylogeny of *Polyplax serrata* lice obtained with Bayesian inference (MrBayes). The values at the nodes show statistical supports (Bayesian posterior probability). Abbreviations of clades: N - nonspecific clade; S - specific clade; W - western lineage of specific clade; E - eastern lineage of specific clade; A—*Apodemus agrarius* and *uralensis* clade.

**Supplementary Figure 4:**
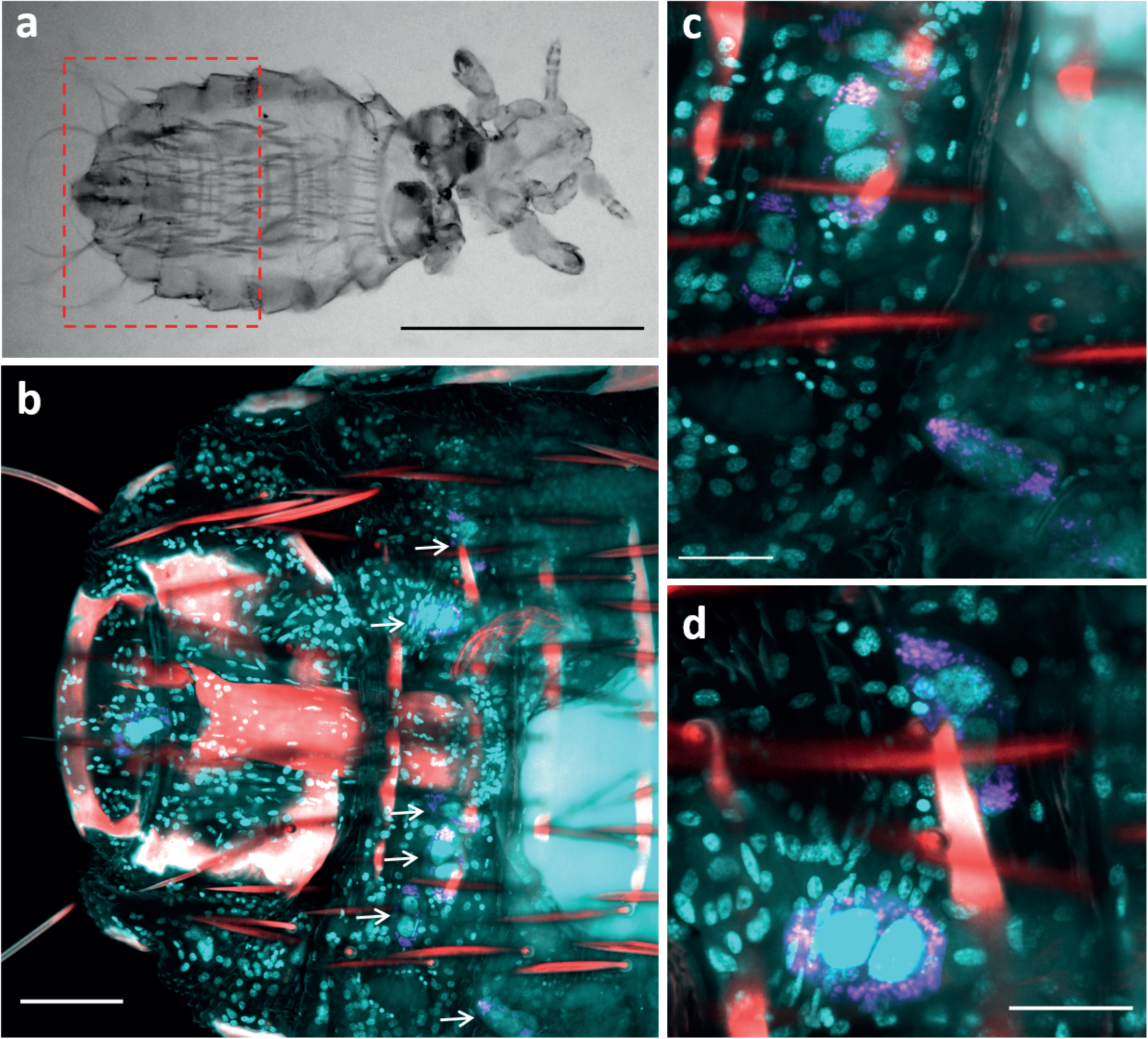
Confocal microscopy of a FISH-stained male *H. acanthopus.* (a) Transmitted light differential interference contrast image showing the louse cut in correspondence of the thorax to allow penetration of paraformaldehyde. The dashed red square defines the region shown in panel b. (b, c and d) Hybridization signals for the generic bacterial probe EUB338 (magenta) show bacterial localization within the potential bacteriocytes. The white arrows in panel b indicate the bacteriocytes shown in panel c and d. Signals for DAPI (cells nuclei in cyan) and tissue autofluorescence (hair shaft and chitin structures in red) were also captured. Scale bars: (a) 500 μm, (b) 50 μm, (c, d) 20 μm.

